# A selfish genetic element and its suppressor causes abnormalities to testes in a fly

**DOI:** 10.1101/2023.02.06.527273

**Authors:** Sophie Lyth, Andri Manser, Tom A. R. Price, Gregory D. D. Hurst, Rudi L. Verspoor

**Affiliations:** Institute of Infection, Veterinary and Ecological Sciences, University of Liverpool, Biosciences Building, Liverpool, UK; Department of Evolutionary Biology and Environmental Studies, University of Zurich, Zurich, Switzerland

**Keywords:** Meiotic drive, X chromosome, gonads, fertility, suppression, *Drosophila subobscura*

## Abstract

Selfish genetic elements (SGEs), specifically X-chromosome meiotic drive (XCMD), create huge conflicts within a host’s genome and can have profound effects on fertility. Suppressors are a common evolutionary response to XCMD to negate its costs. However, whether suppressors themselves can cause negative non-target effects remains understudied. Here, we examine whether the intragenomic conflicts created by XCMD and its suppressor affect gonad morphology in *Drosophila subobscura.* We found significant differences in testes, seminal vesicle, and accessory gland size depending on whether a male carried a non-driving X chromosome, an XCMD, and if the XCMD was suppressed. We also found the first evidence of abnormal testes development that is specifically associated with a suppressor of XCMD. Unlike other studies, our evidence suggests that XCMD in *D. subobscura* creates major abnormalities to male gonads. These abnormalities are most frequent if both XCMD and its suppressor are both present. While costs of suppression have importance in theoretical models, they have largely been ignored in empirical XCMD systems. Overall, this study highlights that genetic conflict, created by SGEs and their suppressors, is a potent evolutionary force that can have major impacts on gonad development and gametogenesis.

## Introduction

Gamete production, and with this testes and ovary development, are shaped by both natural and sexual selection. For instance, egg size is strongly affected by natural selection (1,2) whereas testis size and sperm number is heavily influenced by sexual selection (3,4), with trade-offs between investment in testes and somatic function. Thus, testes are classic examples of organs that have to trade-off between sexual and natural selection (5). However, gonads are also the site of an additional set of trade-offs; the need to defend against selfish genetic elements (SGEs) (6–8).

Challenging the laws of ‘fair’ Mendelian inheritance, SGEs can bias their own transmission into the next generation without increasing the fitness of the carrier and often at the cost of the rest of the genome (9). For example, B chromosomes are additional to an organism’s normal chromosome complement, that carry superfluous DNA (10,11). They accumulate in the ova rather than polar bodies, allowing them to selfishly accumulate in the germline (10). As a result, they are widely reported in nature and can cause phenotypic differences and reduce fertility (10,12,13). Co-evolutionary responses to intragenomic conflicts created by SGEs in gonads are likely to shape their evolution, alongside natural and sexual selection. Thus, the role of SGEs on gonad evolution should not be overlooked, and could be significant and universal because SGEs are both diverse and near ubiquitous in nature (9).

Conflicts created by SGEs can have enormous consequences on the host’s fertility. For instance, *P* element dysgenesis is a property of transposable elements active in *Drosophila simulans* germlines (14). Strong evidence for the impact of SGEs in evolution comes from observations when SGEs enter a naïve host that it has not coevolved with. The *P* element in its native host is not associated with any gross defects (15). Yet when this element is horizontally transmitted into naïve hosts, it becomes associated with a number of abnormalities, including sterility and dysgenic ovaries (16). Another example is X chromosome meiotic drive (XCMD), where male carriers have their Y-sperm killed or damaged thus ensuring the transmission of only the selfish X chromosome to the next generation (17). This reduces a male’s fertility and sperm competitive ability (18–20). Thus, a range of SGEs can negatively impact their hosts and are capable of damaging the gonads and fertility of both sexes. When faced with a multitude of conflicts caused by SGEs, how do organisms and in particular their gonads, respond to deal with these challenges?

Here we investigate how the development of the testes responds when faced with XCMD and a suppressor of this system. XCMD is a type of SGE that creates huge conflicts within the testes (21). XCMD manipulates the process of meiosis in males, where selfish X chromosomes kill Y-bearing sperm during spermatogenesis, resulting in the overproduction of daughters (17). The transmission advantage gained by XCMD can allow it to rapidly spread through populations, distorting sex ratios and even leading to population extinction owing to a lack of males (22–24). Thus, testes in XCMD carrying males are a good model to investigate how SGEs impact the evolution of gonads. Additionally, testes are particularly vulnerable to SGEs due to the asymmetry of gamete production between males and females. Eggs are large and very costly for a female to produce, so anything that damages eggs will be extremely costly and very quickly selected against. On the other hand, because male gametes are generally massively over-produced, damage to sperm may be more readily be tolerated. In cases where males only produce a reduced amount of sperm, they can still be successful, making sperm and testes a good target for the activity of SGEs.

In XCMD, male carriers lose up to half of all the sperm that they produce (25). Despite the lack of sperm produced, drive carrying males do not often show fertility costs in single matings when compared to wildtype males (18,26–29). Indeed, in stalk eyed flies, the presence of sperm-killing XCMD appears to have driven the evolution of a compensatory increase in the testes size of XCMD males (30–32). This allows them to produce a similar amount of sperm to non-XCMD males. These studies, that look at how the testes respond to drive, raises the question as to whether drive systems in general promote changes in the testes.

Meiotic drive is deleterious to male function, and selection commonly favours unlinked suppressor factors that restore Mendelian segregation (21,33). Suppressors, either Y-linked or autosomal, must have the ability to prevent some killing of Y-bearing sperm when a driving X chromosome is present (34). Once a suppressor has evolved, and as long as it has no associated costs, it is expected to rapidly spread to fixation (35,36). Despite this, the impact of suppressors of XCMD on the testes themselves has been rarely considered in empirical studies, despite being recognised as potentially important in theoretical studies (37). Ongoing co-evolution between drive and its suppressor could have different outcomes depending on how a suppressor works, whether it has associated costs, and if it has non-target effects. Thus, the question remains: does the emergence of suppressors negate any testes phenotype (and associated selection) produced by SGEs? Alternatively, the need for testes to evolve could be enhanced if they have to deal with both XCMD and suppression, which are locked in a spiralling co-evolutionary arms race (37–40).

In this paper, we use the XCMD system in *Drosophila subobscura* to look for any effects on testes morphology. *D. subobscura* harbours an XCMD system called ‘SR’ (41). The strength of drive is strong with SR carrying males producing broods that are 85-100% female (41,42). Given this strong drive, a simple expectation is that SR should rapidly spread to fixation everywhere. However, it does not. Instead, in North Africa, SR has remained at a stable frequency of approximately 20% for at least 50 years (41–43). Additionally, *D. subobscura* are reported to be strictly monandrous (28,44) so effects of sperm competition are irrelevant in this species (18,45). We recently discovered suppressors of the SR system (43) that are capable of fully restoring a normal 50:50 sex ratio to SR carrying males. Despite the prediction that suppressors against drive should rapidly fix within populations, this suppressor remains rare at a frequency of 1-5%. The suppressor is also found in multiple locations across North Africa at these low frequencies. The lack of spread of this suppressor within populations may indicate that it is associated with its own costs and creates its own burden upon its host. While we refer to “the suppressor”, we have not fully characterised it. Initial crosses of XCMD into suppressor genotypes results in partial suppression, and several generations of introgression are required to reach complete drive suppression (Supplementary figure 1). Thus, we believe “the suppressor” involves multiple loci across at least two chromosomes, and the Y is not involved. However, in the interests of brevity, we will refer to it as “the suppressor” here, and aim to dissect the complexity in future work.

We determine whether testes morphology is affected when males carry the selfish SR X chromosome (henceforth referred to as SR males) in comparison to standard (ST) males that do not carry SR. We also examine if SR males that also carry the suppressor genes (^Supp^SR males) can restore intact testes or whether the suppressor itself can have its own influence on testes morphology. We used independent SR from multiple locations in Morocco to investigate whether drive in populations have evolved and responded in the same way when it is paired with suppressors that originated from populations in Tunisia.

## Materials and Methods

### Origin and maintenance of fly stocks

Isolines of *Drosophila subobscura* were originally collected from Tabarka, Tunisia (36.95°N 8.74°E) in 2013 (46), Ras el Oud, Morocco (34.15°N 4.00°W), Demnat, Morocco (31.73°N 7.01°W) and Ouzoud, Morocco (32.02°N 6.72°W). Moroccan collections took place in 2019 and all flies were caught using banana, yeast and beer baited traps (47). All isolines were kept in standard *Drosophila* vials on an ASG double yeast medium (10g agar, 85g sucrose, 40g yeast extract, 60g maize, 1000ml H_2_O, 25ml 10% nipagin) at 18°C on a 12:12 light/dark cycle. Experimental flies were generated by collecting unmated flies from isolines from each population. All eclosed flies were removed from populations 24 hours prior to ensure flies had not mated. Fly collection was done using CO_2_ anaesthesia to separate flies by sex. Flies were stored in new vials on a sugar yeast medium (100g sugar, 100g yeast extract, 20g agar, 1000ml H_2_O, 25ml 10% nipagin, 5ml propionic acid) with no more than 10 flies per vial.

### Isolation and introgression of SR X chromosomes

We isolated an SR chromosome from a wild caught fly from each of Ras el Oud, Morocco, Demnat, Morocco and Ouzoud, Morocco, where SR occurs at a frequency of approximately 20% (43). SR males were initially identified though a phenotypic scan by counting their offspring sex ratio. Males that produced more than 85% females were deemed SR males and were used to isolate their driving X chromosomes. In order to produce the SR males (that carry an SR chromosome), suppressed SR males (^Supp^SR), that carry a suppressed version of the SR chromosome, and standard (ST) males (that do not carry the SR chromosome) that were used in this investigation, we introgressed both vulnerable and suppressing ST isolines onto the isolated SR chromosomes. This introgression process meant we were able to produce flies that only differ genetically in carrying drive (males either carried an SR X chromosome or an ST X chromosome) and either carried the suppression genes or genes susceptible to drive. Preliminary experiments have shown that the Moroccan isolines show no suppression against SR (43), thus these isolines were used to produce the SR males that produce 100% daughters. Additionally, the ST isoline from Tunisia contains genes that are capable of suppressing SR. This Tunisian ST isoline was therefore used to produce the ^Supp^SR males that produce a normal 50:50 sex ratio despite the male carrying SR (Supplementary figure 1).

We introgressed the SR X chromosomes onto either the vulnerable Moroccan ST isolines or the suppressing Tunisian ST isoline for 14 to 17 generations (Supplementary figure 2). The introgression process was done in two crosses per generation. Cross 1: females homozygous for the isolated SR X chromosome (‘SRSR’ females) were crossed to a standard (‘ST’) male from the target isoline. This cross produces heterozygous females (‘SRST’ females) that are discarded, and ‘SR’ males that carry the SR X chromosome and the target isoline autosomes. Cross 2: simultaneously, SRSR females were crossed to SR males containing the isolated SR X chromosome. This cross produces only SRSR females. SR males containing the target autosomes generated from cross 1 are then crossed to the SRSR females generated from cross 2 to initiate the subsequent generation. ST males are also re-crossed to the SRSR females from cross 2 to reset cross 1 for the next generation. Over multiple generations the target autosomes will be introgressed onto the SR X chromosomes (Supplementary figure 3).

### Dissection of male testes

We examined the size of male testes and seminal vesicles (SV’s) of SR males, ^Supp^SR males and ST males. To look for changes to male testes that may develop over time as they mature or alternatively be present when just sexually mature and disappear with age, we dissected males when they were either just sexually mature (at 3-days-old) or when they had reached full sexual maturity (at 7-days-old) (48). When at either 3-or 7-days-old, males were anesthetised using FlyNap (Carolina Biological Supply Co., Burlington, USA) and the testes were dissected out of the males in 300µl of phosphate buffered saline (PBS). Once dissected, the testes were transferred onto a microscope slide containing 80µl of PBS. A photograph of the testes was taken under constant light conditions with a microscope mounted Nikon D5100 camera at x40 magnification. To control for body size effects, the left wing of each male was removed using forceps under a microscope (alternatively, the right wing was taken if the left was damaged) and mounted onto a glass microscope slide. A photograph of each wing was taken using the microscope mounted camera.

### Measuring abnormalities to testes

We first examined photographs for evidence of abnormalities in the testes. Only abnormalities in the testes themselves was considered. Abnormality was scored as a binary variable by the presence of physical holes in the orange sheath that coats the testes (Figure 1a). An intact orange sheath that showed thinning was not classed as abnormal. To avoid bias and ensure consistency, this process was double blinded and each photo was independently evaluated by three people. Here, the proportion of males that showed abnormal testes differed by observer. However, when repeating the analysis just using the data of each observer, we found the same effect that ^Supp^SR males experienced more abnormalities (Analysis 1: GLM: F_8,210_ = 11.227, *p* <0.001; Analysis 2: GLM: F_8,210_ = 9.821, *p* <0.001; Analysis 3: GLM: F_8,210_ = 8.818, *p* <0.001) (Supplementary Figure 4). Thus, a composite of the three analyses was used to determine a final testis abnormality score for each male in which testes were assigned as abnormal if two or more evaluators scored them as abnormal.

**Figure 1:**
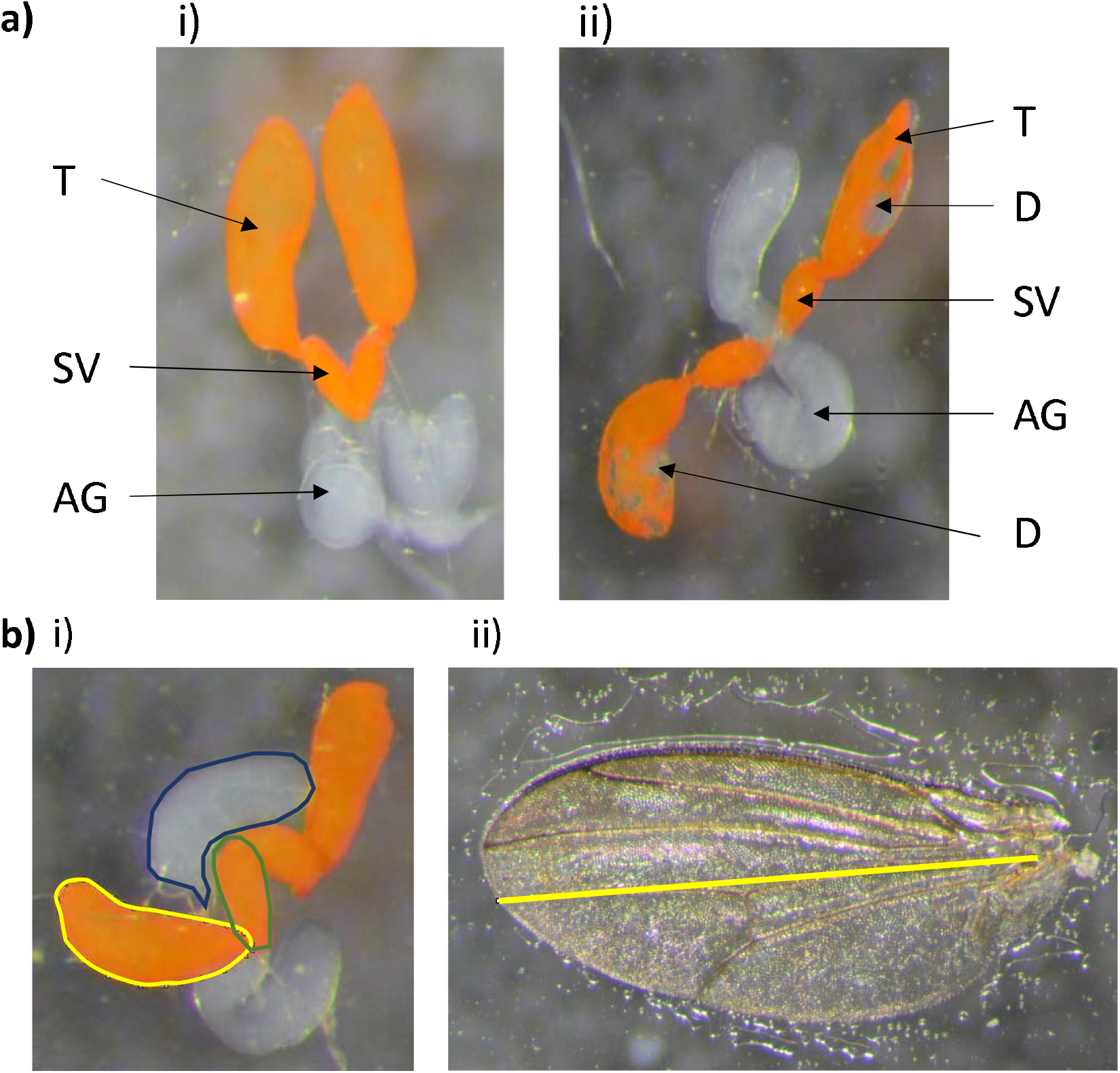
Analysis and measurement of male reproductive organs and wing size. a) Example images of testes (T), seminal vesicles (SV), accessory glands (AG) and visible abnormalities to testes (D). ai) Image of intact testes with no abnormalities to the orange sheath. aii) Abnormal testes defined by the presence of physical holes in the orange sheath. bi) Measurement of testes (yellow), seminal vesicles (green) and accessory gland (blue) area using ImageJ. bii) Measurement of wing length in ImageJ as defined in Gilchrist *et al.,* 2001 (50).

### Measuring testes size, SV size, accessory gland size and wing length

Testes and SV size was measured from images using ImageJ 1.48 (49). A subset of images with visible accessory glands were also analysed in ImageJ to measure accessory gland size. The area of each individual testis was measured (Figure 1b) and a mean of the two taken for each male. If only one testis could be measured for size, it was this single measurement that represented a male’s mean testes size. The same procedure was performed for the SV’s and accessory glands. Wing lengths were also measured in ImageJ to control for body size (Figure 1b), using the method in Gilchrist *et al.,* 2001 (50). The mean testes, SV and accessory gland size for each male was divided by wing length to generate testes, SV or accessory gland size relative to wing length. These relative measurements were used in all statistical analyses.

### Statistical analysis

All statistical analysis was performed using the statistical software R, version 3.6.0 (51). A binomial generalised linear model (GLM) was used to investigate differences in the frequency of abnormalities to male testes. Testes size, SV size and accessory gland size were analysed by fitting a GLM with a gaussian error distribution. All models considered the effects of the type of male (SR, ^Supp^SR, ST susceptible or ST suppressed males), and the age of the male when it was dissected (3-or 7-days old). ST males were split into either ST susceptible or ST suppressed to allow comparisons of the suppressor in the absence of the selfish X chromosome. For testes abnormalities, testes size, and SV size analysis, the original location of origin of the SR X chromosome was included in the model as this provided a better model fit. In these analyses, the effect of SR X chromosome location was significant but idiosyncratic and does not affect the main effects of drive (for individual line effects, see supplementary figures 7-10). For all GLMs, a fully saturated model was produced and was systematically reduced to the minimum adequate model through a step-wise simplification process using Akaike information criterion (AIC). Difference in responses to all GLMs under different treatments were assessed by analysis of variance (ANOVA) followed by a TukeyHSD comparison. For simplicity, all graphs presented align SR next to ST susceptible and ^Supp^SR next to ST suppressed as they share genetic backgrounds to make interpretation clearer of what happens when SR is added to each genetic background.

## Results

### Wing size

Wing size (a proxy for body size), was analysed using a GLM. The model considered the type of male and the age of the male and their interaction. The interaction was not significant and was dropped from the model. Wing size was affected by age (GLM: F_1,218_ = 7.518, *p* = 0.007), where 7-day-old males were smaller than 3-day-old males (TukeyHSD, *p =* 0.007). The type of male also had a significant effect on wing size (Supplementary figure 6: GLM: F_3, 218_ =32.762, *p* < 0.001). ST suppressed males were smaller than ^Supp^SR (TukeyHSD, *p* = 0.002), SR (TukeyHSD, *p* <0.001) and ST susceptible males (TukeyHSD, *p* < 0.001). ^Supp^SR males were smaller than SR and ST susceptible males (TukeyHSD, *p* < 0.001 and *p* = 0.002, respectively). There was no difference in the wing size of SR and ST susceptible males (TukeyHSD, *p* = 0.120).

### Testes size

Mean testes size relative to wing length was analysed using a GLM. The model considered the type of male and the age of the male. A significant age by type of male interaction was found (Figure 2a: GLM: F_3, 213_ =3.663, *p* = 0.0132). Here, ST males (both suppressed and susceptible) have significantly reduced testes size as they age from 3-days-old to 7-days-old (TukeyHSD, *p* < 0.001 for both ST types). This reduction in testes size was not seen in SR and ^Supp^SR males as they aged (TukeyHSD, *p* = 0.975, TukeyHSD, *p* = 0.780, respectively). There was an overall effect of age on testes size (GLM: F_1,213_ = 15.705, *p* < 0.001), where 7-day old-males had larger testes (TukeyHSD, *p* <0.001).

**Figure 2:**
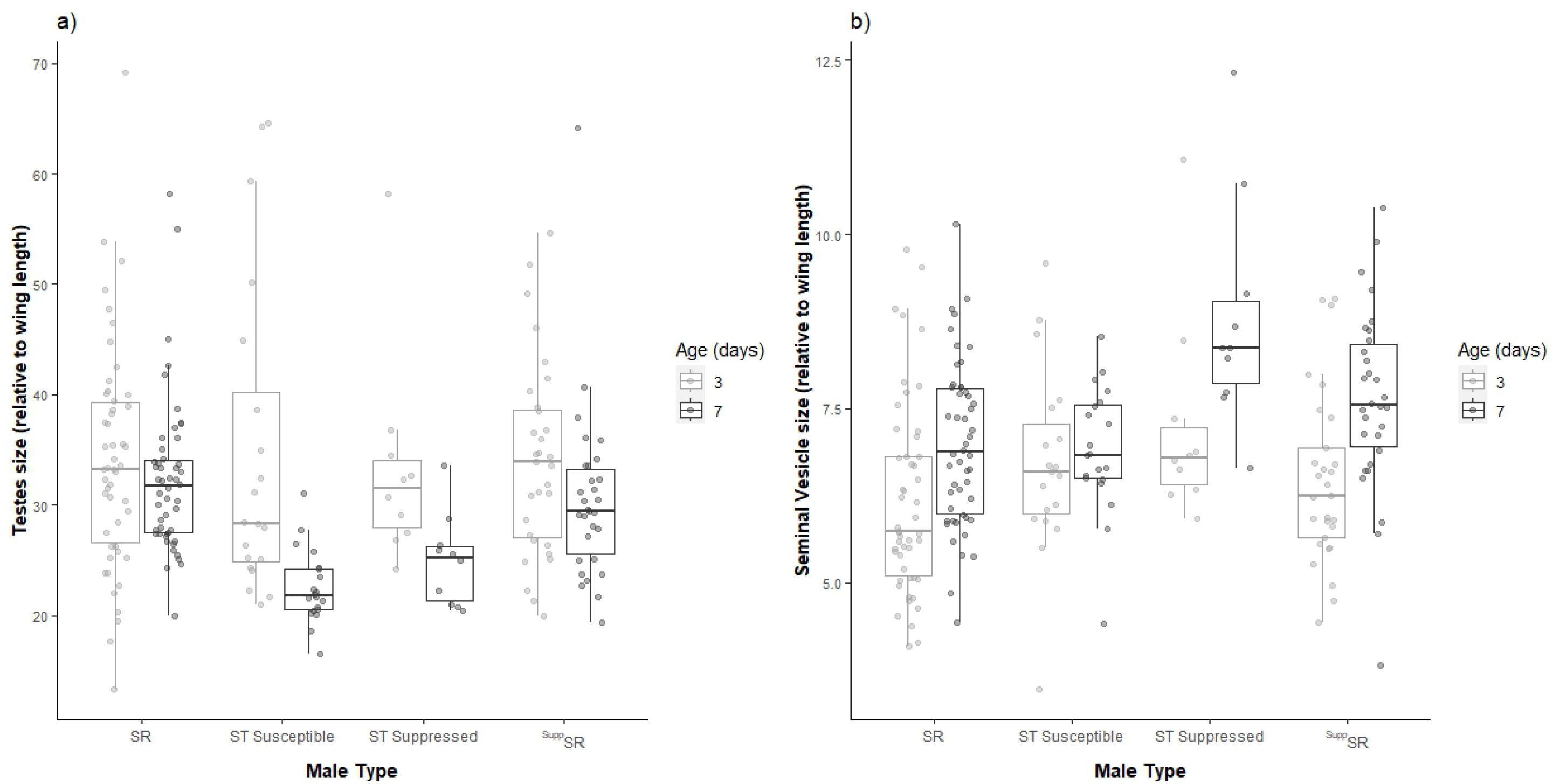
The mean size of a male’s two testes (a) or seminal vesicles (b) of Drosophila subobscura males that were either SR, ST susceptible, ST suppressed or ^Supp^SR. All measurements were relative to individual body size. The boxplots display the upper and lower quartiles, the median and the range. Individual points represent each measurement obtained.

### Seminal vesicle size

Mean SV size relative to wing length was analysed using a GLM. The model considered the type of male and the age of the male and their interaction. The interaction was not significant, so was dropped from the model. Overall, SV’s were bigger in 7-day-old males than 3-day-old males (Figure 2b: GLM: F_1, 213_ = 24.095, *p* < 0.001). The type of the male also had a significant effect on mean SV size (Figure 3b: GLM: F_3, 213_ = 7.384, *p <* 0.001). Here, ST suppressed males have larger SV’s than SR males (TukeyHSD, *p <* 0.01), ST susceptible males (TukeyHSD, *p* = 0.005) and ^Supp^SR males (TukeyHSD, *p* = 0.026). However, there was no difference in SV size between ^Supp^SR and SR and ST susceptible males.

**Figure 3:**
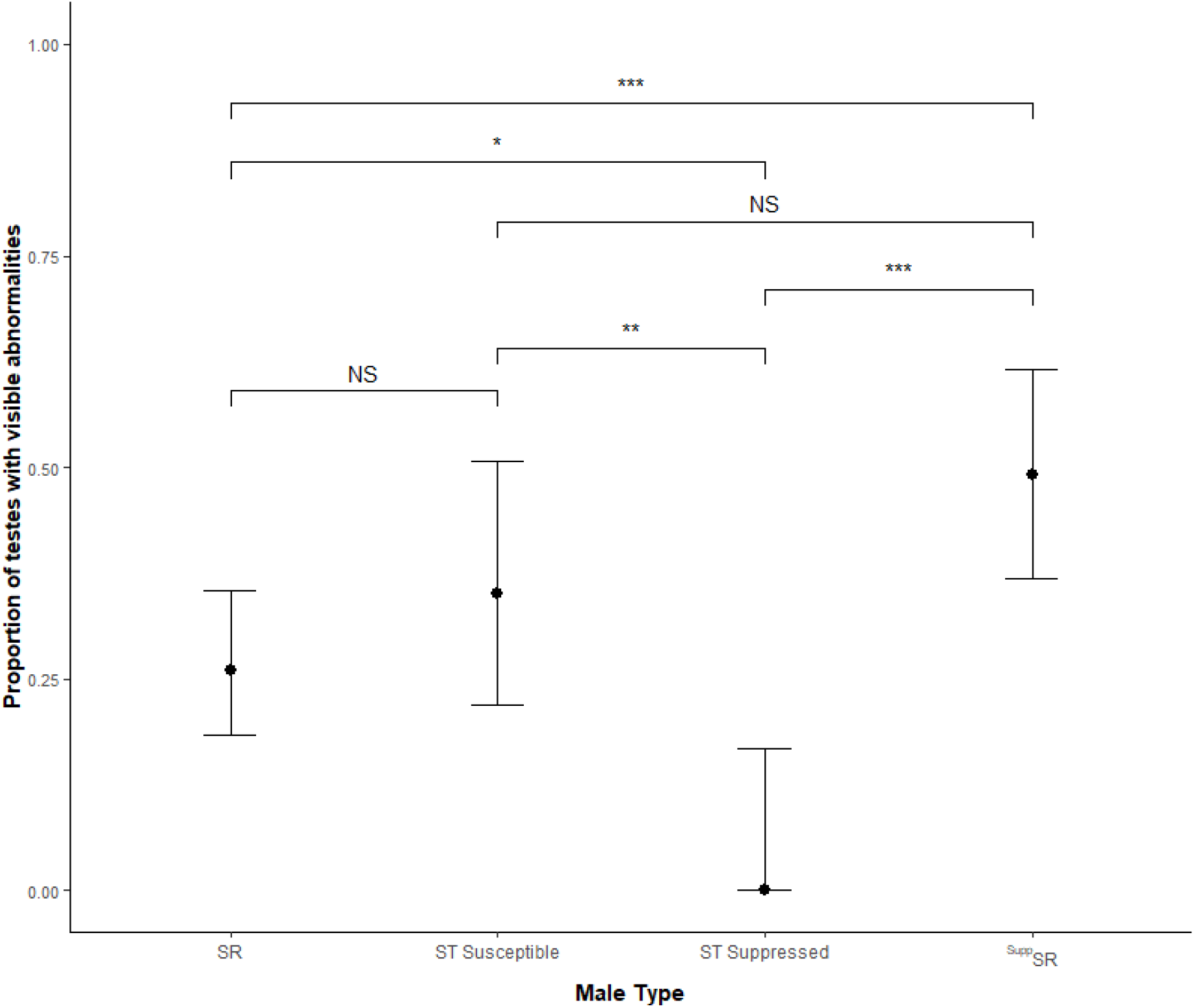
The mean proportion of males that have visible abnormalities to their testes that were either SR males, ST susceptible males, ST suppressed males or ^Supp^SR males, with 95% confidence intervals. The figure also reports statistical comparisons where NS is non-significant, * is *p* < 0.05, ** is *p* < 0.01 and *** is *p* < 0.001.

### Abnormal testes

The presence of visible abnormalities to testes was analysed using a binomial GLM. The model considered the type of male and the age of the male and their interaction. Neither age nor the age by male type interaction had an effect on visible abnormalities to a male’s testes (GLM: F_1, 199_ = 0.064, *p =* 0.801, F_10,199_ = 1.609, *p* = 0.106, respectively) so they were dropped from the model. Male type had a significant effect on the likelihood of male having abnormal testes (Figure 3: GLM: F_3, 210_ = 9.580, *p* < 0.001). While most categories differed from one another, the most noticeable differences were that ^Supp^SR males had a significantly higher proportion of abnormal testes compared to ST suppressed males (where by our measure, 49% of ^Supp^SR males had visibly abnormal testes in comparison to 1% ST suppressed males) (Figure 3: TukeyHSD, *p* < 0.001). On the other hand, SR males did not differ significantly in the frequency of abnormal testes in comparison to ST susceptible (18% of SR males had damage to their testes in comparison to 22% in ST susceptible males, TukeyHSD, *p* = 0.594). ^Supp^SR also had a higher amount of damage than SR males (TukeyHSD, *p* = 0.0015). An example of both normal and abnormal testes can be seen in Figure 1a.

We also dissected the testes from SR, ^Supp^SR and ST susceptible males when 7-days-old for preliminary and exploratory analysis of the internal structure of each male type’s testes (see supplementary figure 11 for details). SR and ^Supp^SR testes showed disorganisation, with gaps between cells, while ST susceptible males showed normal tightly packed sperm development (supplementary figure 11). While ^Supp^SR males showed a higher level of organisation than SR males, the internal structure of male testes did not resemble that of ST susceptible males.

### Accessory gland size

Accessory glands were measured in any images where they were clearly visible. Mean accessory gland size relative to wing length was analysed using a GLM. The model considered the type of male and the age of the male. Models analysing accessory gland size did not consider the individual location the SR X chromosome originated from due to the smaller sample size and having little power to detect differences. A significant age by type of male interaction was found (Figure 4: GLM: F = 3.689, *p* = 0.0135). Here, ^Supp^SR males’ accessory glands increase in size as they age (TukeyHSD, *p* = 0.004). This change in accessory gland size was not seen in SR males (TukeyHSD, *p* = 0.999), ST susceptible males (TukeyHSD, *p* = 0.999) or ST suppressed males (TukeyHSD, *p* = 0.933). Male type also had an effect on accessory gland size (GLM: F = 3.689, *p* = 0.0135). Overall, ^Supp^SR males had larger accessory glands than SR and ST susceptible and ST suppressed males (TukeyHSD, *p* = 0.046, TukeyHSD, *p* < 0.001, TukeyHSD, *p* = 0.037, respectively). SR males also had larger accessory glands than ST susceptible males (TukeyHSD, *p* = 0.003). Finally, the age of the male had an effect on accessory gland size (GLM: F_1, 143_ = 5.713, *p* = 0.0181). Overall, 7-day-old males had larger accessory glands than 3-day-old males (TukeyHSD, *p* = 0.019).

**Figure 4:**
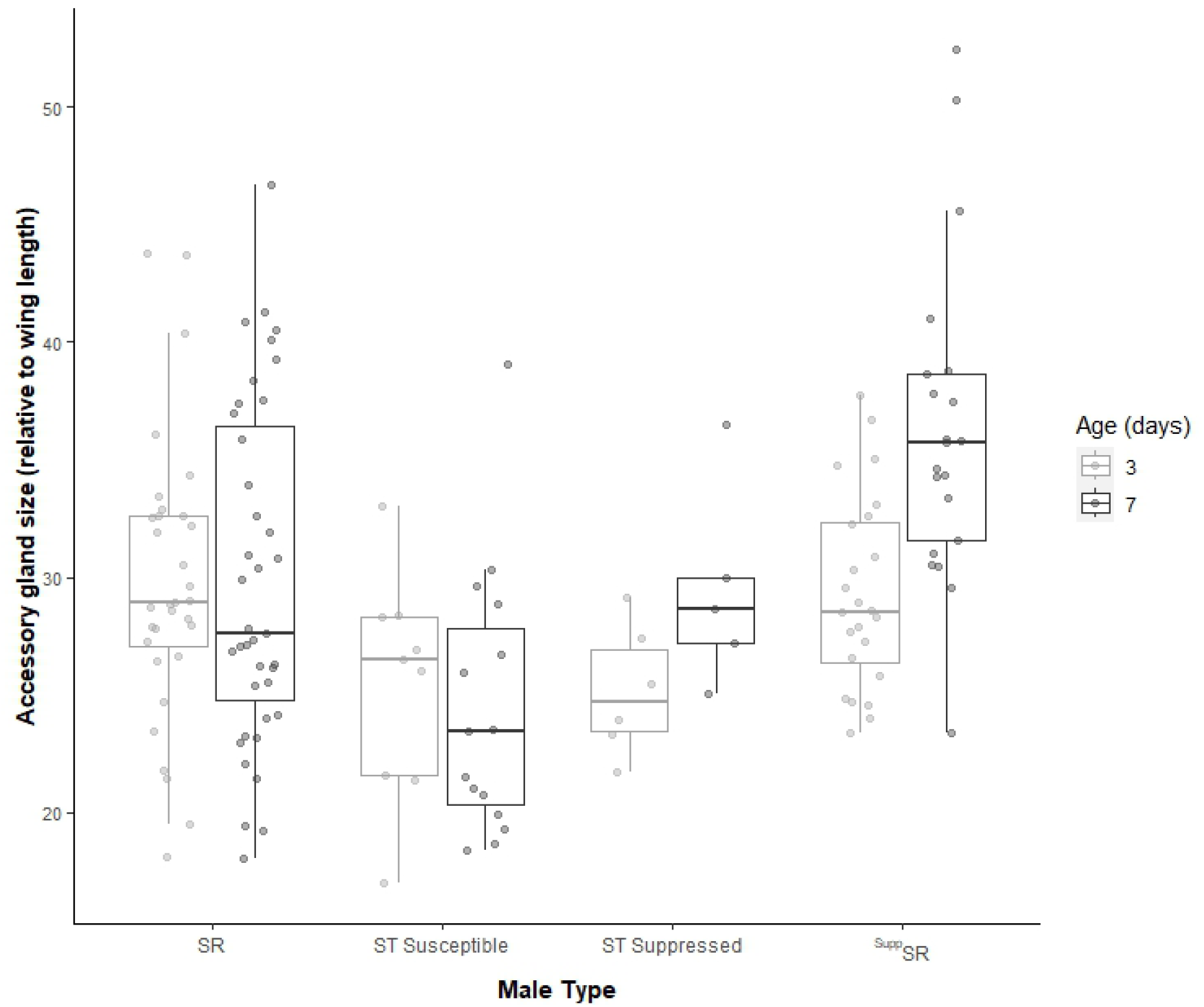
The mean size of the two accessory glands of male *Drosophila subobscura* that were either SR, ST susceptible, ST suppressed or ^Supp^SR. All measurements were relative to individual body size. The boxplots display the upper and lower quartiles, the median and the range. Individual points represent each measurement obtained.

## Discussion

XCMD and their suppressors generate potent genetic conflict that may also be a force influencing gonad development, morphology and evolution (31,32). In this study, we examined whether the intragenomic conflicts created by SR and its suppressor affects gonad morphology in *D. subobscura.* We found significant differences in testes, seminal vesicle, and accessory gland size depending on whether a male carried XCMD or not, and also if XCMD was suppressed. We also found the first evidence of extreme whole-organ abnormal development of testes associated with carrying a suppressor of XCMD. While males with SR from independent origins had differences in the morphological traits observed, this variation was idiosyncratic and overall the main effect of SR and its suppressor were the same, suggesting these effects are robust across populations. In conclusion, we found strong evidence that XCMD and its suppressor can have a major impact on gonad development and morphology of its carriers.

### Testes and Seminal Vesicle size

We found evidence that the testes of ST (both suppressed and susceptible) males, but not those of SR and ^Supp^SR males, reduce in size as they age. *Drosophila subobscura* are strictly monandrous (28,44), and males may not benefit from high investment in continually producing large quantities of sperm (52–54). In particular, if *D. subobscura* can partition their ejaculates, like in numerous species across a wide array of taxa (55,56), they may only transfer to a female the sperm she will need. This results in reduced pressure on the testes to continually produce sperm (and the costs of spermatogenesis this entails). In contrast to the reduction in testes size observed in ST males, it appears that SR and ^Supp^SR testes remain the same size as males age. We propose three mutually non-exclusive reasons for this: (1) compensation, (2) damage or (3) a delay in sperm production.

In stalk-eyed flies, XCMD males have enlarged testes to compensate drive induced sperm mortality, and males produce an equivalent number of sperm to normal males (31). This compensation of male testes appears to occur from an early development stage (32). Compensation may also be the effect that we have observed in *D. subobscura*. Indeed, a previous study has suggested that the loss of Y-bearing sperm in SR males is compensated through cyst doubling in the testes (57,58). Unlike in stalk-eyed flies, we do not observe this increase in testes size at all ages. At 3-days-old, all male types have equivalent sized testes. If this was compensation, would SR male carriers not always have enlarged testes? In addition, in a species with no sperm competition (28,44), loss of sperm number may not need to be compensated.

Contrary to the compensation hypothesis, damage may instead be inflicted upon the testes resulting in their enlarged size. The internal testes structure of SR and ^Supp^SR males showed disorganisation, with gaps between cells, while ST testes showed normal tightly packed sperm development (supplementary figure 11). This may indicate that XCMD carriers instead experience internal testes damage and/or spermatogenic failure, the severity of which may accumulate over time. Indeed, the gapping between cells and their general disorganisation may explain why SR and ^Supp^SR male have enlarged testes at this age. However, a full investigation of the internal morphology of testes and associated sperm development is required to explore this further. Currently, little is known about the interaction of XCMD and genes generally involved in gonad development (59,60).

Finally, if testes are largest simply when the process of spermatogenesis is at its peak, then sperm production, and the reaching of a full sperm complement, may simply be delayed in SR and ^Supp^SR males. As a result, testes would remain larger for a longer period of time, while they continue full sperm production to an older age. A detailed temporal breakdown of sperm production and storage would be required to confirm this hypothesis.

We found no strong effects of carrying XCMD on SV size, as these were equivalent in all male types, other than ST suppressed. The reasons for this are unclear and do not provide strong support for any of our above theories. SVs are sperm storage organs in *Drosophila* males, where mature sperm that develop in the testes move into the SV for storage until mating (61,62). Thus, in unmated males, SV size is generally related to the amount of sperm stored (63). The equivalent SV size may indicate that these male types are all producing and storing a similar amount of sperm, providing some additional support for the compensation theory. Conversely, all the sperm stored in the SVs of SR and ^Supp^SR males may not be viable. It is possible that these males are bundling dead or damaged sperm that is incapable of fertilisation or zygote formation. The disruption to the internal structure of these testes suggests that spermatogenesis is likely to be affected in these males (supplementary figure 11). A full examination of the viability of sperm stored within the SVs would be required to confirm this.

### Whole testes abnormalities

Remarkably, we found evidence that, when the suppressor is paired with a driving SR X chromosome, it causes gross visible abnormalities to testes. Overall, these abnormalities are less in XCMD males that lack the suppressor which is comparable to that of ST susceptible males. Once evolved, suppressors against drive are usually expected to remove or minimise the negative impacts of drive (33,64,65). However, we demonstrate for the first time that suppression, while mitigating some negative effects of XCMD, can also have its own effects on testes morphology, that likely imposes costs. In addition, given the scope of the morphological differences we see, suppression genes may influence general testes development that, when expressed and paired with SR, results in their abnormal development and potential spermatogenesis failures. In addition, these abnormalities are not restricted to the external testes. Our preliminary SEM analysis of the internal structure of male’s testes shows that both SR and ^Supp^SR males have a high level of disorganisation and gapping between cells, while ST shows normal tightly packed sperm development. Thus, SR appears to be having its own influence of testes morphology, that becomes enhanced when the suppressor is also present. This is in support of our theory that SR and its suppressor are inflicting damage or developmental changes upon a male’s testes. To what effect, if any, these abnormalities then has on a male’s fertility remains an open question.

It is critical these abnormalities, and any associated costs is understood as it could affect the balance between drivers and suppressor that may provide insight into the long-term stability of drive that we see in many natural systems (43,66–68). Additionally, if suppressors can spread despite deleterious effects, they have the potential to fix negative mutations within a population, which is rarely considered in empirical studies. There are many cases of cryptic (or muted) drive systems that are fixed within populations (64,65,69). Here, drive systems are only revealed through outcrosses with other populations or species that do not have the suppressor. These factors are especially important to consider in relation to artificial drive systems (70). What would happen to artificial drive if a suppressor unexpectedly evolved and fixed deleterious mutations within the target population? The evidence we have found that major abnormalities are associated with suppression against XCMD suggests that not only can SGEs carry huge costs, but in addition suppressors and their interactions with SGEs must be accounted for.

### Accessory gland size

The extent of changes to male gonads as a result of XCMD seem not to just be limited to the testes. We observed changes to the whole of the male gonads, including the accessory glands. Accessory glands are organs that are responsible for the production of seminal fluid that contains accessory gland proteins (71). In *Drosophila*, accessory gland proteins influence female behaviour and a males sperm competitive ability (72,73). In some systems, accessory gland size is also correlated with male mating rate and success (74,75). In this study, we found that ^Supp^SR accessory glands are the largest, followed by SR which were larger than ST (both suppressed and susceptible) males. Carrying SR may be associated with compensatory increases in investment in accessory gland proteins to ensure their mating success rather than transferring an equivalent amount of sperm to ST males. Additionally, the highest investment in accessory glands was from ^Supp^SR. When SR males are suppressed they also endured the highest frequency of testes abnormalities, which may indicate males are investing more in gaining a mate. Despite this, it remains unclear as to what the role of accessory glands is in a monogamous species where the importance of them becomes largely unknown.

### Conclusion

Here, we demonstrate that the presence of XCMD and its suppressor can cause significant morphological changes to male gonads. This is the second system, that we are aware of, where XCMD strongly influences male gonad development (31,32). Unlike the XCMD system in stalk-eyed flies, where morphological changes were deemed a compensation mechanism against drive, our evidence suggests that XCMD in *D. subobscura* inflicts gross abnormalities on male gonads that becomes even more severe in the presence of its suppressor. This highlights that SGEs are a strong evolutionary force that can broadly impact gonad development and gametogenesis. Additionally, considering drive alone may only reveal half of the story. Suppressors can also have their own severe effects on their carriers. Overall, the fitness effects of either an SGE or its suppressors may be severe for individuals that carry them.

## Acknowledgements

The authors would like to thank Jola Tanianis for her help with the isolation of the SR chromosomes, and Kieran Romain for measuring the accessory glands. The authors acknowledge the use of the biomedical electron microscopy unit provided by Liverpool shared research facilities, Faculty of Health and Life Sciences, University of Liverpool. We would also like to thank the three anonymous referees for their helpful comments on the manuscript. This work was funded by the National Environment Research Council ACCE DTP and the grant no. NE/S001050/1.

