## Supplementary material for "A selfish genetic element and its suppressor causes abnormalities to testes in a fly"

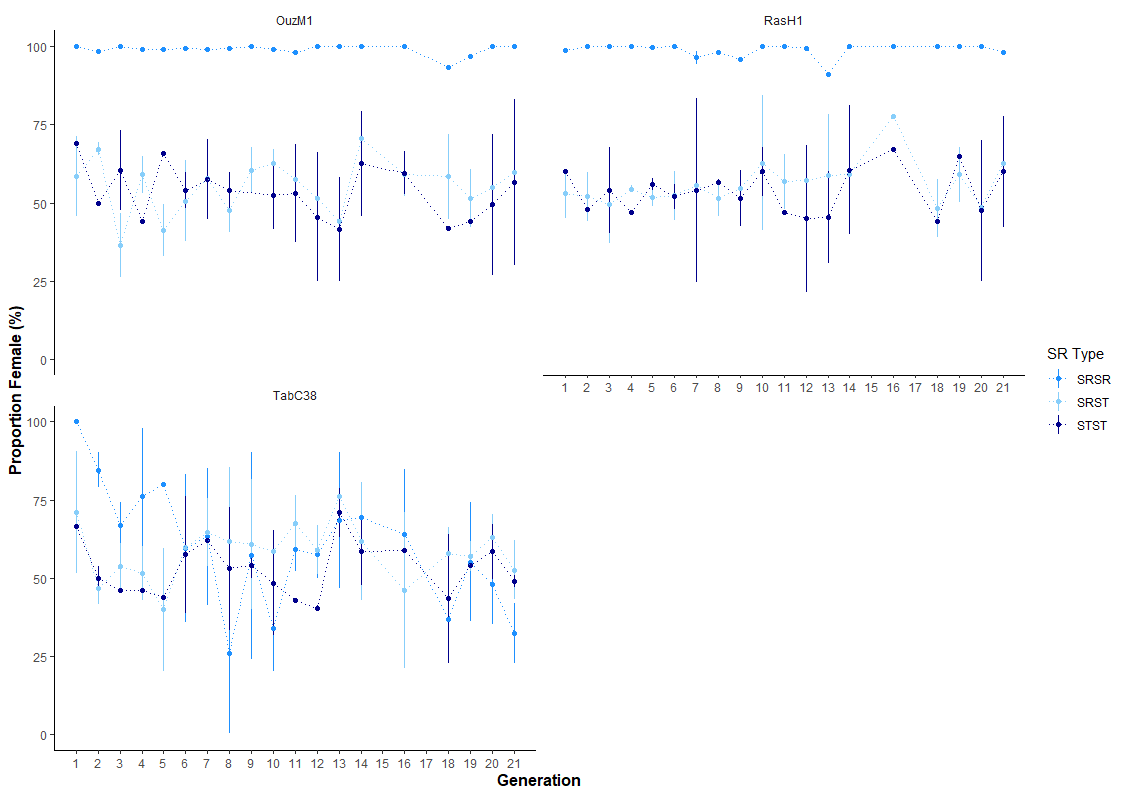
**Supplementary figure 1:** Sex ratio counts of each generation of the introgression. SRST is cross 1 from supplementary figure 3, SRSR is cross 2 and STST is cross 3. OuzM1 and RasH1 are the Moroccan ST lines that are susceptible to SR. This is shown by the SRSR line only producing females. TabC38 is the Tunisian suppressing line shown by males appearing in the SRSR line. This suppression is also complete as the sex ratio in this line is returned to 50:50. Here, the points are the mean proportion female across 5 replicates and the error bars are the 95% confidence intervals. The absence of error bars occurs when the number of replicates that could be counted was less than three.

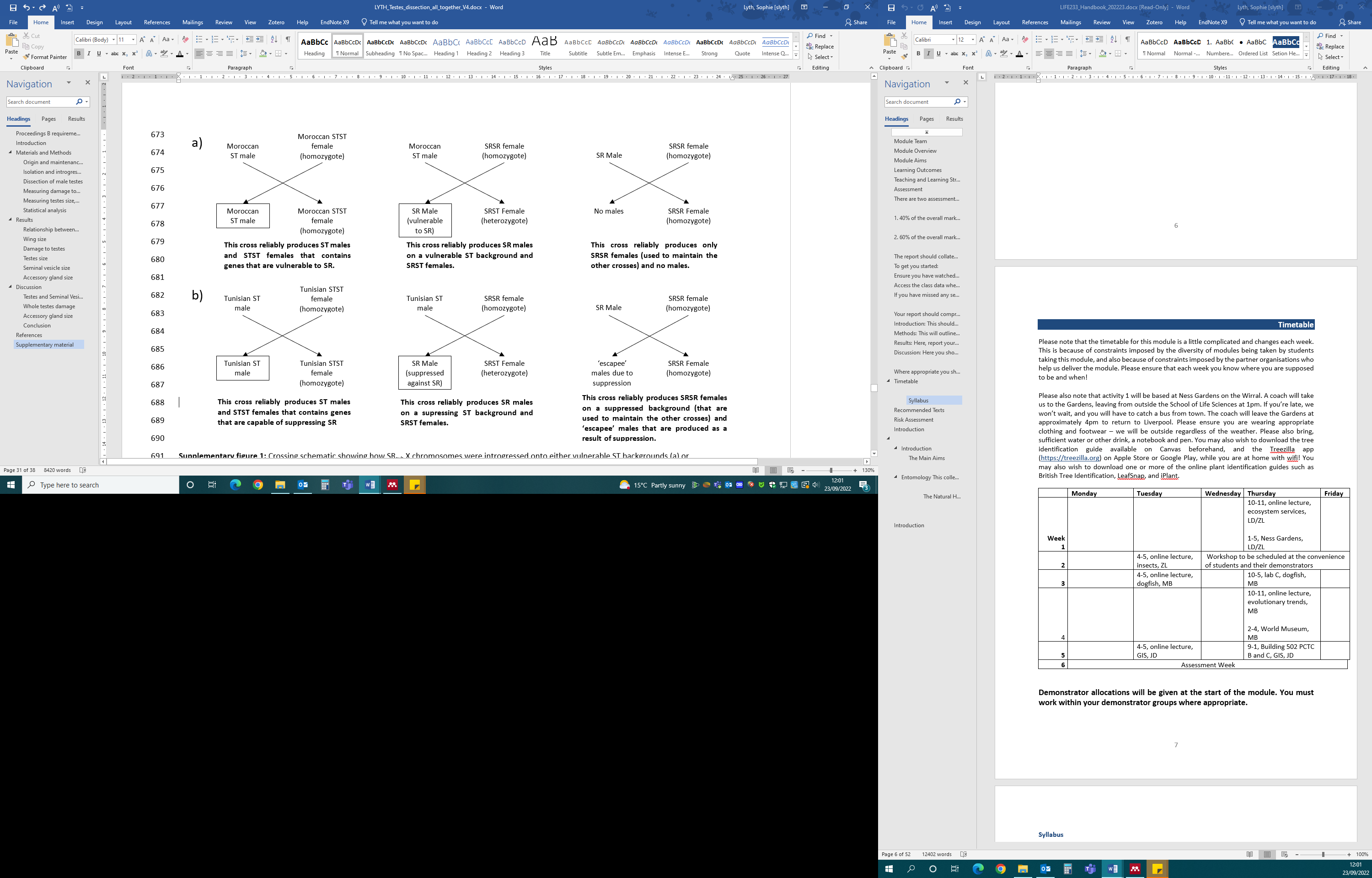

**Supplementary figure 2:** Crossing schematic showing how SR X chromosomes were introgressed onto either vulnerable ST backgrounds (a) or suppressing ST backgrounds (b) to produce either ST males, SR males ^Supp^SR males (outlined) used in this study.

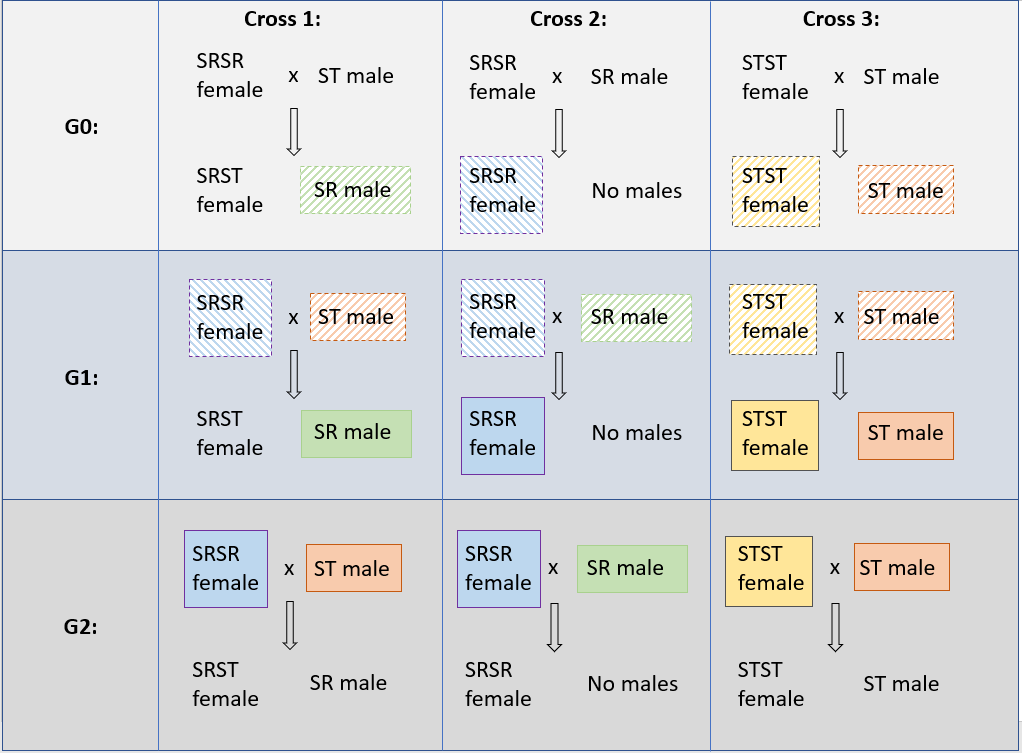
**Supplementary figure 3:** Crossing schematic showing the detailed crosses used for the introgression of SR onto ST autosomal backgrounds. ST lines were either of Moroccan (susceptible to SR) or Tunisian (can suppress SR) descent. Here, G stands for generation. The colour coding of the fly genotypes shows how the emerging flies of one generation’s cross are then used to set the subsequent generation whilst continuing to introgress SR onto the targeted ST autosomal background. For example, the solid green SR males produced in G1 are used as the father in the G2 SRSRxSR cross, and the G0 diagonal blue SRSR females are used as mothers in both the G1 SRSR x ST cross and the G1 SRSR x SR cross.

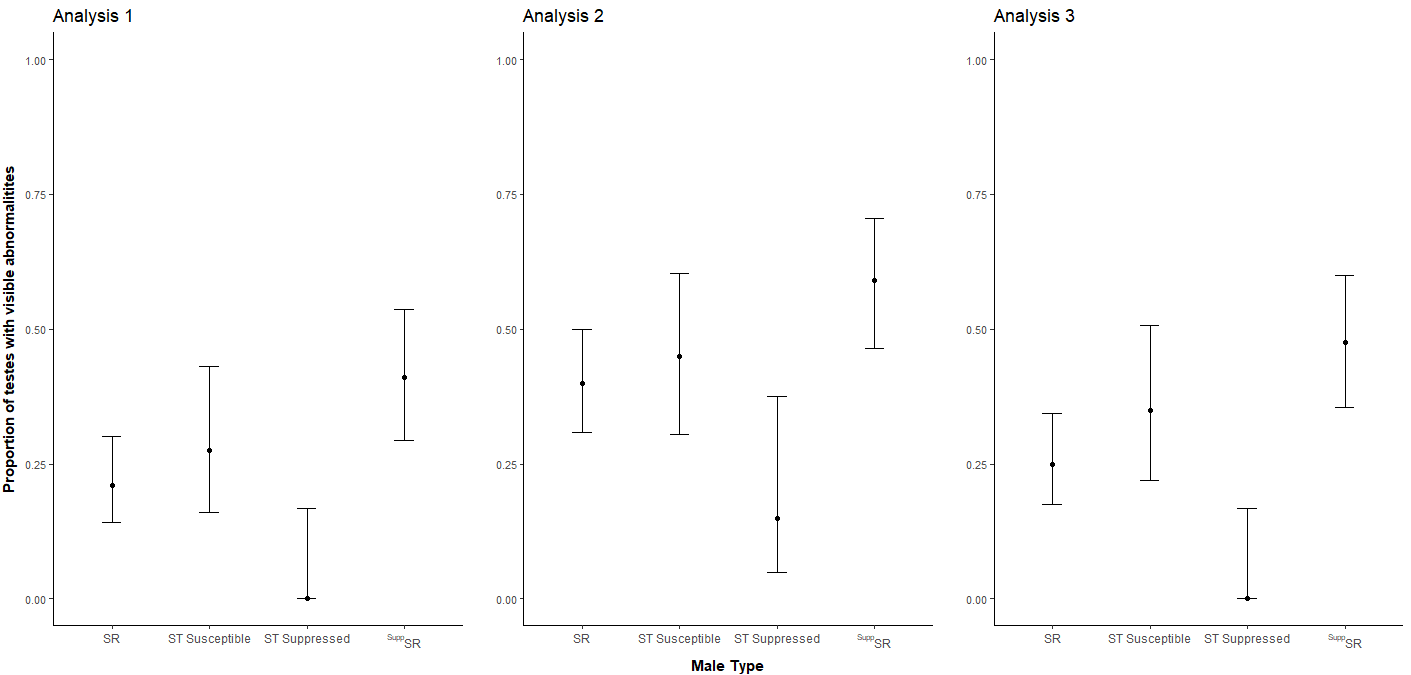
**Supplementary figure 4:** The three independent analyses made by three people of the mean proportion of males that have visible abnormalities to their testes. The photographs were analysed blind and independently three times to avoid bias during analysis. Although the proportion of abnormalities to each type of male changed per analysis, overall all three analyses found there was a higher number of ^Supp^SR that had visible abnormalities to their testes. Here, the points represent the mean proportion of males that had abnormal testes (means presented as back transformed) and the error bars represent 95% confidence intervals.

### **Relationship between wing length and testes, SV or accessory gland size**

Relationships between wing length (a standard proxy measure of body size) and either testes size, SV size or accessory gland size were examined through a Spearman’s rank correlation. There was a positive correlation between wing length and mean testes size (Supplementary figure 5a; ρ = 0.292, n = 221, *p <* 0.001). Here, as body size increased, so did the mean testes size. However, there was no relationship between wing length and mean SV size (Supplementary figure 5b; ρ = 0.002, n = 221, *p* = 0.980) or wing length and accessory gland size (Supplementary figure 5c; ρ = 0.086, n = 151, *p* = 0.292).

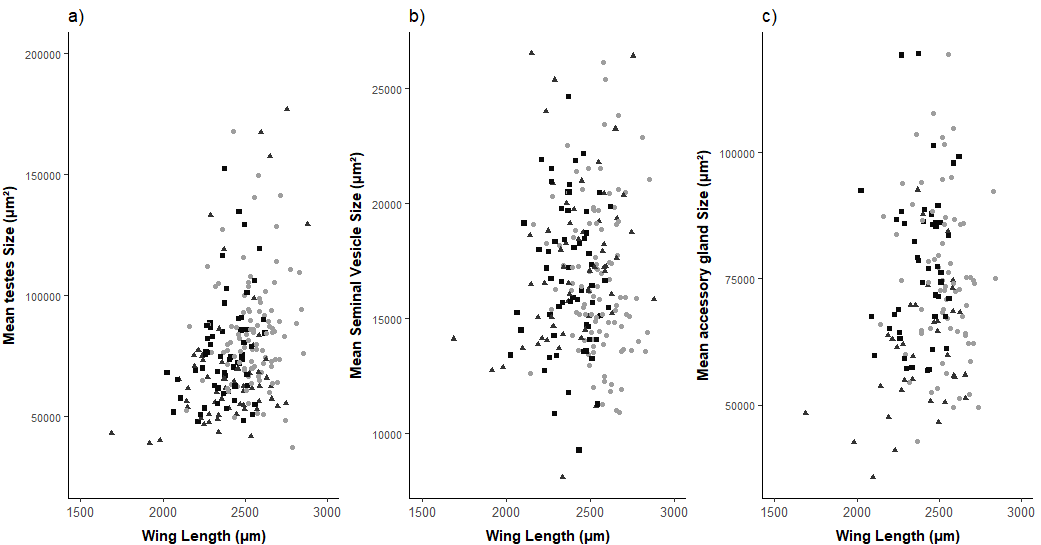

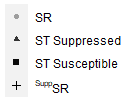

**Supplementary figure 5:** The relationship between wing length (µm) and mean testes size (a), mean seminal vesicle size (b) or mean accessory gland size (c). Each point refers to an individual male that was either an SR male (circle), ST suppressed male (triangle), ST susceptible (square) or ^Supp^SR male (cross).

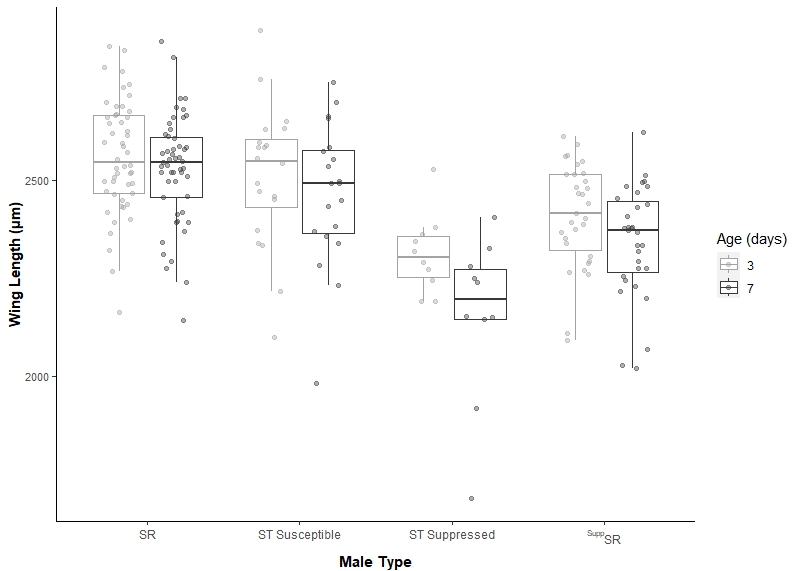
**Supplementary figure 6:** Boxplot of wing length, used as a proxy for body size, of *Drosophila subobscura* males that were either SR, ST susceptible, ST suppressed or ^Supp^SR males. The boxplots display the upper and lower quartiles, the median and the range. Individual points represent each measurement obtained.

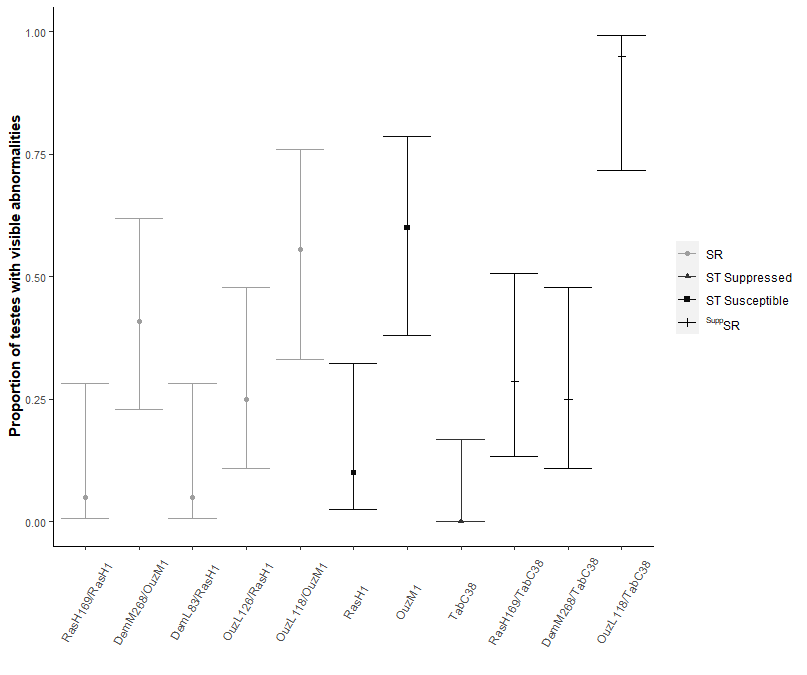
**Supplementary figure 7:** The mean proportion of males that have visible abnormalities to their testes, split by location of SR origin. Here, the points represent the mean proportion of males that has abnormal testes(means presented as back transformed) and the error bars represent 95% confidence intervals.

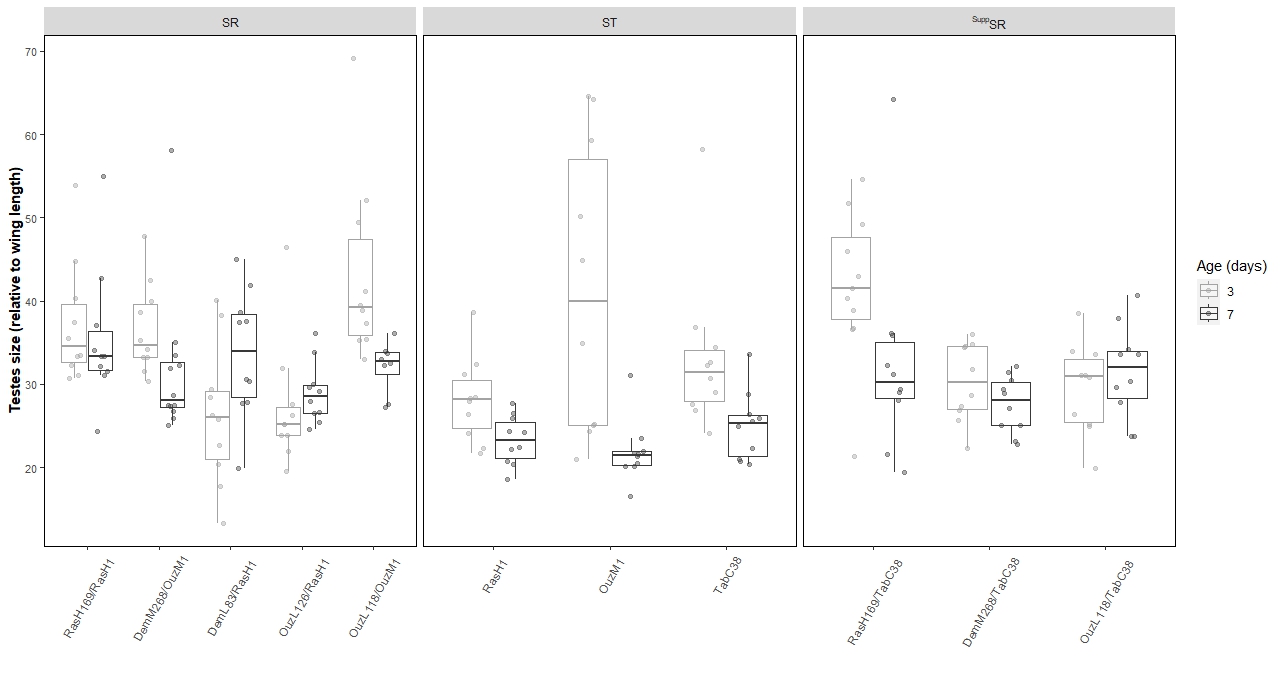
**Supplementary figure 8:** Mean testes size of *Drosophila subobscura* males relative to body size split by location of SR X chromosome origin.

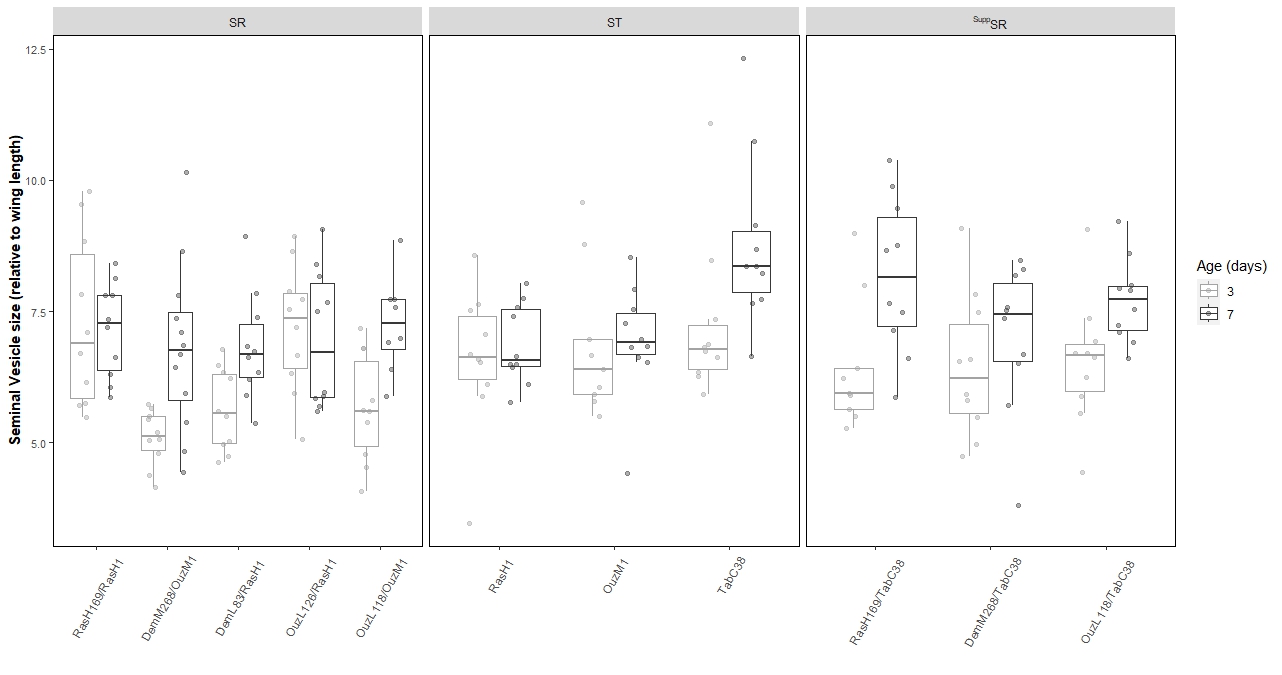
**Supplementary figure 9:** Mean seminal vesicle size of *Drosophila subobscura* males relative to body size split by location of SR X chromosome origin.

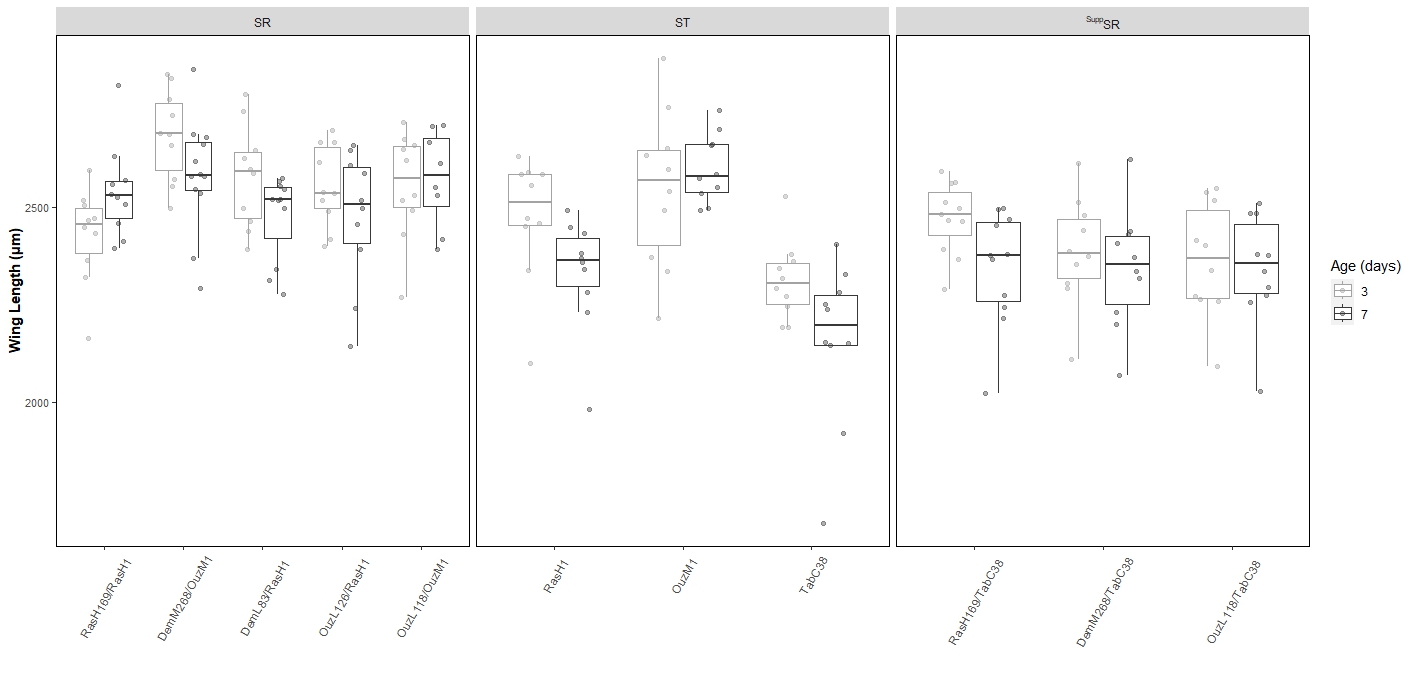
**Supplementary figure 10:** Wing length, used as a proxy for body size, of *Drosophila subobscura* males split by location of SR X chromosome origin.

**Supplementary figure 11:** Serial block-face scanning electron microscopy (SBFSEM) images of (a) ST susceptible, (b) SR and (c) ^Supp^SR male testes when 7-days-old. These images are a preliminary analysis of the internal structure of male testes. Image a) shows normal testes structure, with tightly packed developing sperm bundles, all with organised spiral structure of internal cells. Images b) and c) show general disorganisation, with loose packing of sperm bundles, cells showing irregular alignment (IA) and gapping between cells (G). These images independently support the size differences of the different male’s testes and also highlight differences in the overall shape of the testes when looking at their cross section. For Serial block-face scanning electron microscopy (SBFSEM), testes were fixed in 4% paraformaldehyde and 2.5% (w/v) glutaraldehyde in 0.1 M cacodylate buffer (pH 7.4). Heavy metal staining consisted of reduced osmium (2% (w/v) OsO_4_, 1.5% (w/v) potassium ferrocyanide in ddH_2_O), 1% (w/v) thicarbohydrazide, 2% OsO_4_ (w/v in ddH_2_O), followed by 1% (w/v) aqueous uranyl acetate overnight at 4 °C. Testes were further stained the following day with Walton’s lead aspartate (0.02 M lead nitrate, 0.03 M aspartic acid, pH 5.5) at RT. Fixation and staining steps were performed in a Pelco Biowave®Pro (Ted Pella Inc., Redding, CA) at 100 w 20 Hg, for 3 min and 1 min, respectively with ddH_2_O washes in between. Dehydration was performed in a graded series of ethanol before filtration and embedding in hard premix resin (TAAB, Reading, UK). Images were acquired using Gatan 3View serial block-face system (Gatan, Pleaseanton, CA) installed on a FEI Quanta 250 FEG scanning electron microscope (FEI Company, Hillsboro, OR). Image stacks were taken at various resolutions: Image A - chamber pressure 70Pa, 3.4Kv, pixel size 18.8x18.8x100nm, dwell time 4us. Image B - chamber pressure 70Pa, 3.4Kv, pixel size 30x30x100nm, dwell time 3us. Image C - chamber pressure 70Pa, 4Kv, pixel size 34x34x100nm, dwell time 4us.

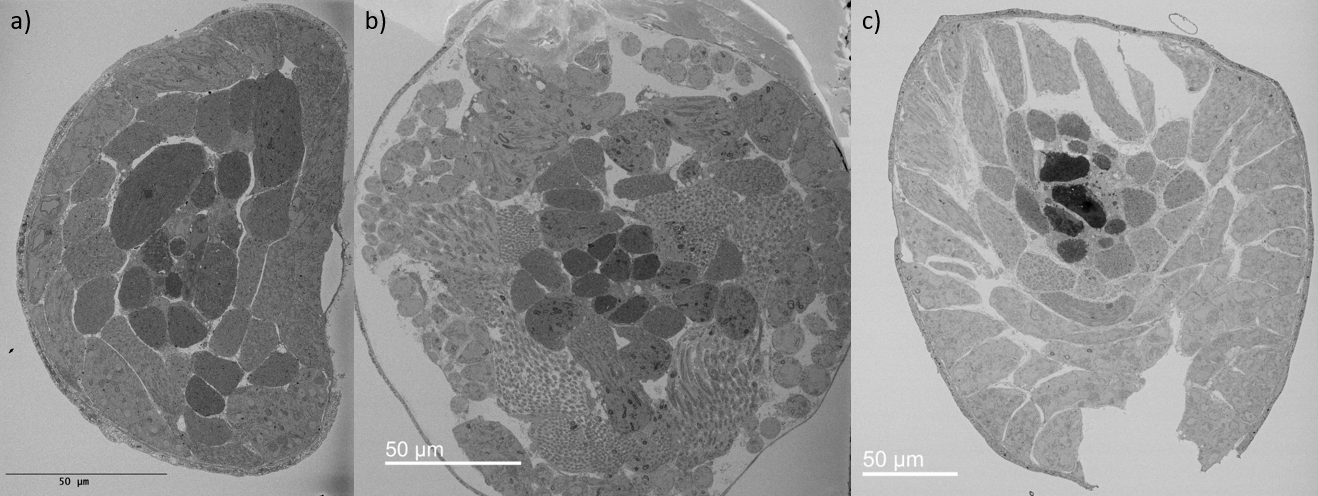

**IA**

**IA**

**G**

**G**
